# Analytical Validation of a COVID-19 qRT-PCR Detection Assay Using a 384-well Format and Three Extraction Methods

**DOI:** 10.1101/2020.04.02.022186

**Authors:** Andrew C. Nelson, Benjamin Auch, Matthew Schomaker, Daryl M. Gohl, Patrick Grady, Darrell Johnson, Robyn Kincaid, Kylene E. Karnuth, Jerry Daniel, Jessica K. Fiege, Elizabeth J. Fay, Tyler Bold, Ryan A. Langlois, Kenneth B. Beckman, Sophia Yohe

**Affiliations:** Department of Laboratory Medicine & Pathology, Division of Molecular Pathology and Genomics; University of Minnesota Genomics Center; M Health Fairview Molecular Diagnostics Laboratory; Department of Genetics, Cell Biology, and Development; Department of Microbiology and Immunology and Center for Immunology; Biochemistry, Molecular Biology and Biophysics Graduate Program; Department of Medicine, Division of Infectious Diseases and International Medicine, University of Minnesota, Minneapolis. 55455.

**Author notes:** Address Correspondence To: Andrew C. Nelson, M.D., Ph.D., Sophia Yohe, M.D.

## Abstract

The COVID-19 global pandemic is an unprecedented health emergency. Insufficient access to testing has hampered effective public health interventions and patient care management in a number of countries. Furthermore, the availability of regulatory-cleared reagents has challenged widespread implementation of testing. We rapidly developed a qRT-PCR SARS-CoV-2 detection assay using a 384-well format and tested its analytic performance across multiple nucleic acid extraction kits. Our data shows robust analytic accuracy on residual clinical biospecimens. Limit of detection sensitivity and specificity was confirmed with currently available commercial reagents. Our methods and results provide valuable information for other high-complexity laboratories seeking to develop effective, local, laboratory-developed procedures with high-throughput capability to detect SARS-CoV-2.

## Introduction

Coronavirus disease 2019 (COVID-19) is a newly emergent pandemic infectious disease caused by Severe Acute Respiratory Syndrome Coronavirus 2 (SARS-CoV-2). SARS-CoV-2 is easily transmitted among humans, has infected at least 649,900 people and caused 30,249 deaths as of March 28, 2020 (Johns Hopkins Coronavirus Resource Center). The need for widespread testing, not just among individuals with symptoms, but health care workers and individuals who have had contact with infected individuals is needed to curb the spread of SARS-CoV-2.

The preferred method for diagnosis of coronavirus infections is by one-step quantitative reverse transcriptase PCR (qRT-PCR) [1-4]. For the majority of qRT-PCR procedures, RNA must be extracted from patient samples [1,3]. SARS-CoV-2 present in patient samples must be inactivated during the lysis step of RNA extraction for the safety of individuals working with the samples. The CDC has confirmed the use of several RNA extraction kits with external lysis buffer that is effective at inactivating SARS-CoV-2 [5]. Unfortunately, access to these kits is severely limited, hampering widespread implementation of testing. Validation of new RNA extraction kits and qRT-PCR reagents is desperately needed to ease supply constraints and to increase testing worldwide. Two of the CDC approved kits, the QIAmp Viral RNA kit (Qiagen) and EasyMag NucliSENS kit (biomérieux), have lysis buffers that contain guanidinium thiocyanate and guanidine thiocyanate, respectively. A third kit, the NucleoSpin Virus RNA/DNA extraction kit (Machery Nagel, Takara) which is not currently CDC approved, contains guanidine hydrochloride in the lysis buffer. We compared these three extraction methods and performed qRT-PCR with RUO primer-probe sets targeting the CDC approved 2019-nCoV_N1, 2019-nCoV_N2 and human RNase P (RP) sequences. Responding to current shortages in reagent availability, we developed a 384-well, lower volume qRT-PCR assay with an alternative single step master mix. Our data demonstrate all three RNA extraction kits can be used with this laboratory-developed procedure to effectively detect SARS-CoV-2 and that reducing input and qRT-PCR reaction volumes can still provide sensitivity of detection down to 5 copies of viral genome per microliter (comparable to previously validated methods).

## Methods

### Sample collection

A synthetic SARS-CoV-2 Standard control (Exact Diagnostics; abbreviated EDx) at 200 cp/µL viral nucleic acid and 75 cp/µL human gDNA was diluted into EDx Negative control (human only) as indicated. The EDx controls are manufactured to require extraction procedures to serve as a synthetic spike-in source for validation of the entire assay procedure and are ddPCR quality controlled for copy number. Synthetic viral RNA sources were acquired from BEI (NR-52358) and ATCC (VR3276T); a genomic isolate of USA-WA1/2020 was acquired from BEI. Plasmid cDNA was acquired from IDT encoding relevant target sequences from SARS-CoV-2 (nCov2, 10006625) as well as SARS (10006624) and MERS (10006623) viruses (to serve as specificity controls) and human RPP30 (RP, 10006626) as a control. Non-identifiable, residual clinical biospecimens were provided by the Minnesota Department of Health (MDH) for use under Common Rule exemption.

### Sample Tracing

Samples were tracked in a Laboratory Information Management System (LIMS) throughout the process from accessioning through to data release. Container barcodes used for tracing did not include any PHI. At initial scanning into the LIMS, each unique sample swab barcode was associated with a unique 0.5-ml 2D-barcoded tube (Micronic), into which the extracted RNA was eluted at completion of extraction. Racks of these 2D barcoded tubes from extraction batches were arranged into 96-well run-plate layouts, which were bottom-scanned to confirm well location into the LIMS, which then generated PCR run plate layouts, ingested the RT-PCR result files for inspection and interpretation by pathologists, and generated report file outputs for import into laboratory information and electronic medical records systems.

### Biosafety Procedures

When handling clinical samples prior to viral inactivation, we use standard practices for biosafety level 2, with enhanced personal protective equipment (BSL2+). This includes a disposable gown, face protection with either full shield or mask with eye protection, Tyvek sleeves, double gloves, and a clean room tacky mat located at the BSL2+ room’s threshold. All work with patient samples is performed in a biological safety cabinet (BSC), including centrifugation, vortexing and heating. Additional BSL2+ precautions include: changing outer gloves after any manipulation of samples prior to removing hands from the BSC, reduced pipette speed to minimize aerosol risks, working over towels soaked in 10% bleach, and disposing pipet tips used with clinical samples into a container with bleach. For viral inactivation, clinical samples are first placed in tubes containing viral lysis buffer and treated according to manufacturer’s protocol. Samples are transferred into a new clean tube containing 96-100% ethanol as per the next step in the RNA extraction protocol. After inactivation, sample tubes are surface decontaminated, placed in secondary containment, and moved into a separate room where the remainder of the RNA extraction protocol is carried out at standard BSL2 practices in a BSC. Personal protective equipment for RNA extraction include gloves, gowns and eye protection.

### RNA extraction

Samples were extracted using three different RNA extraction kits: 1) Qiagen QIAamp Viral RNA Mini kit (abbreviated QIA, catalog # 52906), 2) Macherey-Nagel Nucelospin Virus, Mini kit (abbreviated MN, catalog # 740983.50), and 3) biomérieux easyMag NucliSENS system (abbreviated EMAG). All extraction methods followed manufacturer recommended protocols with the notable exceptions of using 100 µL of starting material and eluting with 100 µL of appropriate elution material as indicated by manufacturer protocols.

### RT-qPCR

#### RT-qPCR setup

Three separate 10 µL RT-qPCR reactions were set up in a 384-well Barcoded plate (Thermo Scientific) for either the N1, N2, or RP primers and probes. 2.5 µL extracted RNA was added to 7.5 µL qPCR mastermix comprised of the following components:

1.55 µL water

5 µL GoTaq^®^ Probe qPCR Master Mix with dUTP (2X) (Promega, Cat # A6120 and A6121)

0.2 µL GoScriptTM RT Mix for 1-Step RT-qPCR (Promega, Cat # A6120 and A6121)

0.75 µL primer/probe sets for either N1, N2, or RP (IDT, Cat# 10006713)

#### Primers and probes

were obtained from IDT (2019-nCoV CDC RUO Kit, 500 rxn (IDT, Cat# 10006713) with the CDC-recommended sequences which can be found at this web address: https://www.cdc.gov/coronavirus/2019-ncov/downloads/rt-pcr-panel-primer-probes.pdf.

#### 20 ul reaction volume comparisons

For 20 µL reaction volume comparison, the reactions were scaled up uniformly from the 10 µL volume. Three separate 20 µL RT-qPCR reactions were set up in a 384-well Barcoded plate (Thermo Scientific) for either the N1, N2, or RP primers and probes. 5 µL extracted RNA was added to 15 µL qPCR mastermix comprised of the following components:

3.1 µL water

10 µL GoTaq^®^ Probe qPCR Master Mix with dUTP (2X) (Promega, Cat # A6120 and A6121)

0.4 µL GoScriptTM RT Mix for 1-Step RT-qPCR (Promega, Cat # A6120 and A6121)

1.5 µL primer/probe sets for either N1, N2, or RP (IDT)

#### qPCR cycling conditions

Reactions were cycled in a QuantStudio QS5 (ThermoFisher) for one cycle of 45°C for 15 minutes, followed by one cycle of 95°C for 2 minutes, followed by 45 cycles of 95°C for 15 seconds and 60°C for 1 minute. A minimum of two no template controls (NTCs) were included on all runs. Baselines were allowed to calculate automatically, and a ΔRn threshold of 0.5 was selected and set uniformly for all runs. Ct values were exported and analyzed in Microsoft Excel. Amplification curves were manually reviewed.

#### 55°C annealing/extension temperature comparisons

To compare the performance of the use of 55°C and 60°C annealing/extension temperatures, reactions were cycled in a QuantStudio QS5 (ThermoFisher) for one cycle of 45°C for 15 minutes, followed by one cycle of 95°C for 2 minutes, followed by 45 cycles of 95°C for 15 seconds and 55°C for 1 minute. We determined that with the reagents and instrumentation used, the 60°C annealing/extension temperature recommended by the mastermix manufacturer (Promega) produced more robust amplification (data not shown).

#### Results Interpretation

|  |  |
| --- | --- |
| N1 AND N2 positive Ct < 40 | Positive for SARS-CoV-2 |
| N1 OR N2 positive Ct < 40 | Inconclusive |
| N1 AND N2 negative; RP Ct < 36 | Negative |
| N1 AND N2 negative; RP Ct > 36 | Invalid (Quantity Not Sufficient) |

## Results

### Preliminary Assay Characterization

An initial pilot experiment aimed to prove acceptable performance of the 384 well assay format with a 10 µL assay volume. We utilized a subset of available clinical biospecimens, synthetic positive extraction controls, and nucleic acid controls to test initial parameters, confirm reproducibility, and determine a initial limit of detection. One aliquot each of four clinical samples (two positive, two negative) and three separate aliquots of a synthetic positive control with 200 viral copies per microliter (EDx_200) were extracted with Qiagen viral RNA reagents (one of the CDC-approved extraction methods); all unique extractions were run in PCR replicate (n=8 clinical replicates, 6 synthetic replicates). All samples showed expected positive (viral N1 and N2 gene targets detected with internal human RP gene control detected) or negative (N1, N2 undetermined, RP detected) results across all replicates. The average Ct values demonstrated a narrow standard deviation across both extraction and PCR replicates (Table 1). Further, we tested five-fold dilutions of input RNA into nuclease free water for all of these samples. Each dilution remained positive for SARS-CoV-2; and demonstrated delta Ct values ranging between 2.01-3.16 for the viral N1 and N2 targets (Table 2). The internal human control RP gene target showed the greatest amount of delta Ct variability in this test.

**Table 1:** Average Ct Values (StdDev) For Synthetic Controls and Pilot Clinical Samples

| Sample | N1<br>Avg. Ct<br>(SD) | N2<br>Avg. Ct<br>(SD) | RP<br>Avg. Ct<br>(SD) |
| --- | --- | --- | --- |
| *EDx_200*<br>*N = 6* | 30.43<br>(0.44) | 30.58<br>(0.29) | 29.59<br>(0.11) |
| *POS5*<br>*N = 2* | 29.14<br>(0.14) | 30.19<br>(0.20) | 28.83<br>(0.19) |
| *POS8*<br>*N = 2* | 18.10<br>(0.11) | 19.36<br>(0.11) | 28.53<br>(0.12) |
| *NEG1*<br>*N = 2* | n/d | n/d | 31.60<br>(0.08) |
| *NEG2*<br>*N = 2* | n/d | n/d | 27.08<br>(0.15) |

N = number of PCR replicates; EDx\_200 was extracted in three replicates

**Table 2:** Average Ct Values for 1:5 Dilutions and Delta Ct vs. Straight Replicates

| SAMPLE | N1 Ct<br>delta Ct | N2 Ct<br>delta Ct | RP Ct<br>delta Ct |
| --- | --- | --- | --- |
| *EDX_200_1:5* | 32.44 | 33.03 | 32.44 |
| *delta Ct* | 2.01 | 2.45 | 2.85 |
| *POS5_1:5* | 32.23 | 33.28 | 32.41 |
| *delta Ct* | 3.09 | 3.09 | 3.58 |
| *POS8_1:5* | 21.26 | 22.22 | 31.8 |
| *delta Ct* | 3.16 | 2.86 | 3.27 |
| *NEG1_1:5* | n/d | n/d | 34.51 |
| *delta Ct* |  |  | 2.91 |
| *NEG2_1:5* | n/d | n/d | 29.82 |
| *delta Ct* |  |  | 2.74 |

Synthetic nucleic acid controls (Integrated DNA Technologies) and extracted RNA from a SARS-CoV-2 clinical isolate (USA-WA1/2020) were also used as direct PCR inputs to demonstrate specificity of the assay. The clinical isolate and plasmid nCov2 controls were both positive for the N1 and N2 reactions and negative for RP, as expected. SARS and MERS nucleic acid specificity controls (run in triplicate) were negative for all three targets, also as expected (Table 3).

**Table 3:** Proportion of Control Replicates Positive for Assay Targets

| Sample | N1 | N2 | RP |
| --- | --- | --- | --- |
| USA-WA1/2020 | 1/1 | 1/1 | 0/1 |
| nCoV_IDT | 1/1 | 1/1 | 0/1 |
| SARS_IDT | 0/3 | 0/3 | 0/3 |
| MERS_IDT | 0/3 | 0/3 | 0/3 |
| RP_IDT | 0/1 | 0/1 | 1/1 |

In this pilot experiment, we performed a initial limit of detection using two separate synthetic RNA SARS-CoV-2 sources (BEI NR-52358 and ATCC VR3276T) with a 2-fold dilution series ranging from approximately 100 viral copies per microliter (cp/µL) to 3 cp/µL. The assay successfully detected both samples down to a calculated lower boundary of 2.8 cp/µL with maximum observed Ct values of 36.19 (Table 4).

**Table 4:** Initial Limit of Detection Using Nucleic Acid Controls

| Sample | Input<br>cp/μL | N1 Ct | N2 Ct |
| --- | --- | --- | --- |
| NR-52358 | 100 | 28.10 | 28.59 |
|  | 50 | 29.39 | 29.83 |
|  | 25 | 30.29 | 30.69 |
|  | 12.5 | 31.38 | 31.71 |
|  | 6.25 | 32.12 | 32.87 |
|  | 3.215 | 33.06 | 33.82 |
| VR3276T | 90 | 31.33 | 31.44 |
|  | 45 | 32.17 | 32.79 |
|  | 22.5 | 32.78 | 33.17 |
|  | 11.25 | 34.26 | 34.09 |
|  | 5.625 | 35.00 | 36.11 |
|  | 2.8125 | 35.06 | 36.19 |

Our preliminary experiments also included parallel assessment of a 20 µL assay format mimicking the volume setup of the CDC-specified assay. We set up PCR replicates for a subset of samples on both volume formats to compare average Cts and coefficients of variation (CV) between the volume formats. The key objective was to determine if the decreased sample input (2.5 µL) of our 384-well format assay significantly compromised assay performance in comparison to the larger volume (5 µL sample input) format. We observed small shifts in raw Ct values of <0.1 to 0.5 for the N1 and N2 targets (Table 5). We transformed replicate Ct values to relative quantity (copy number) to calculate the true quantitative CV across data points for each sample/target on the two assay formats (Table 6). The copy number CVs for clinical samples ranged from 6-14%; the synthetic extraction standard CVs (at 200 cp/µL) ranged from 6-21%.

**Table 5:** Average Ct Values of PCR Replicates Comparing 10 and 20 µL Assay Formats

| Sample | N1 |  | N2 |  | RP |  |
| --- | --- | --- | --- | --- | --- | --- |
|  | Avg_Ct_10 | Avg_Ct_20 | Avg_Ct_10 | Avg_Ct_20 | Avg_Ct_10 | Avg_Ct_20 |
| EDx_200 | 30.4 | 30.1 | 30.5 | 30.2 | 29.5 | 29.5 |
| POS5 | 29.1 | 29.1 | 30.2 | 29.9 | 28.8 | 28.9 |
| POS8 | 18.1 | 18.1 | 19.4 | 18.9 | 28.5 | 28.6 |
| NEG1 | n/a | n/a | n/a | n/a | 31.6 | 31.8 |
| NEG2 | n/a | n/a | n/a | n/a | 27.1 | 27.3 |

**Table 6:**
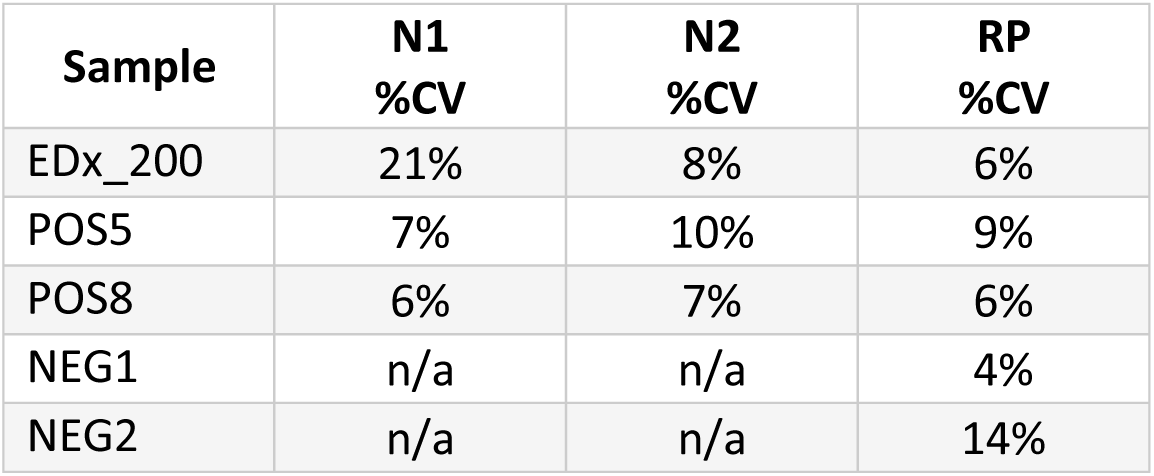
Percentage Coefficient of Variation between 10 and 20 µL Assay Format

| Sample | N1<br>%CV | N2<br>%CV | RP<br>%CV |
| --- | --- | --- | --- |
| EDx_200 | 21% | 8% | 6% |
| POS5 | 7% | 10% | 9% |
| POS8 | 6% | 7% | 6% |
| NEG1 | n/a | n/a | 4% |
| NEG2 | n/a | n/a | 14% |

Quality review of the amplification curves throughout the preliminary assay characterization experiments was performed to optimize the ΔRn threshold value. We noted that artifactual priming events in early amplification cycles would occasionally produce non-exponential amplification traces (Figure 1). A subset of these non-specific traces could cross if auto-thresholding set by the instrument was low (ΔRn threshold 0.2-0.3). We noted that all preliminary true positive data points, including the initial limit of detection experiment, demonstrated robust exponential amplification curves which crossed a 0.5 ΔRn threshold prior to 40 cycles; no artifactual traces reached this cutoff. Therefore we set this threshold for all downstream validation experiments.

**Figure 1:**
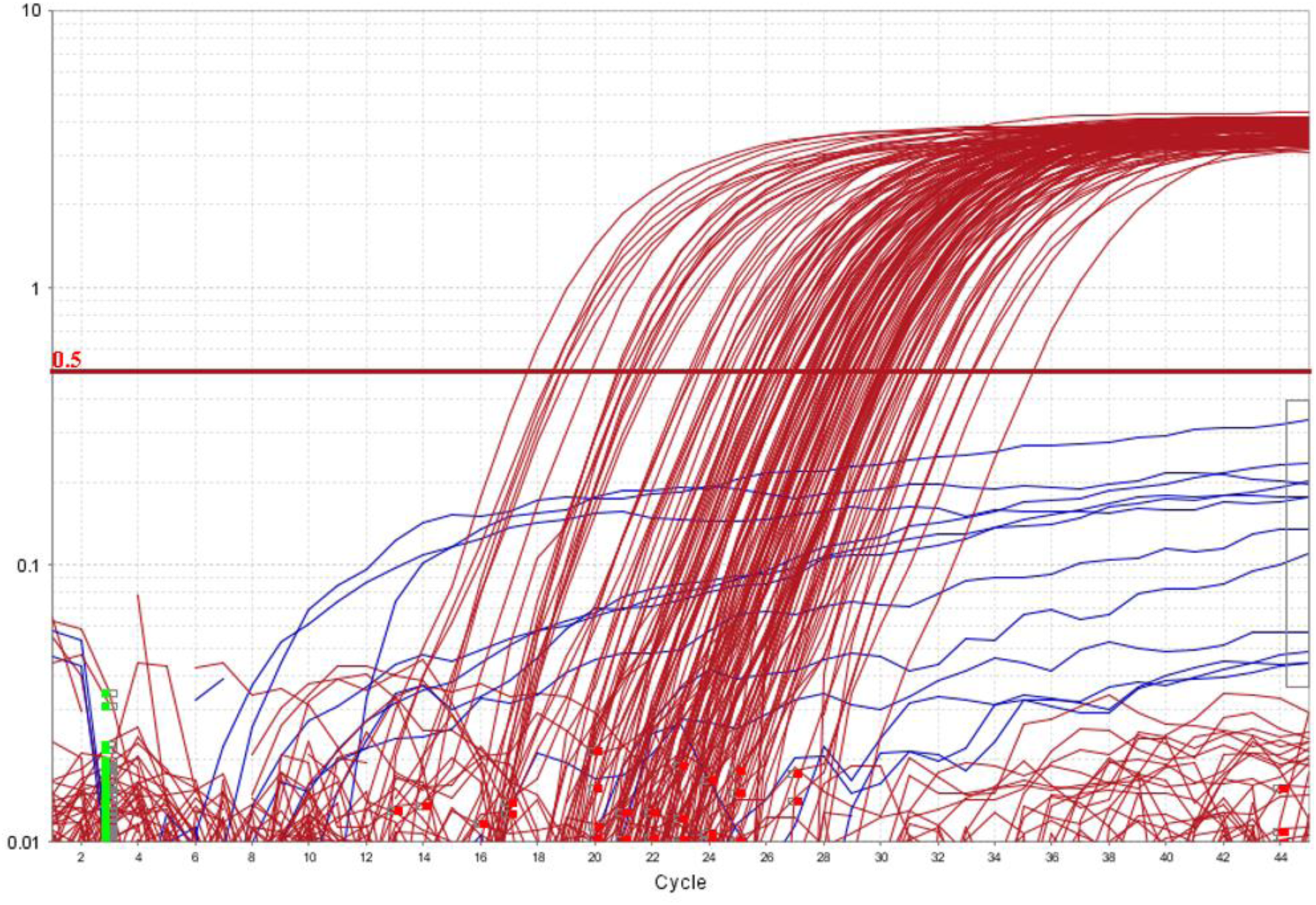
Representative amplification curves of known negative samples. Exponential amplification curves for the RP internal control target are shown in red. Non-specific, non-exponential curves for either N1 or N2 targets are shown in blue. ΔRn threshold was set at 0.5 to protect against potential false positive calls.

### Analytical Accuracy on Clinical Biospecimens

Analytical accuracy evaluation was then performed on ten positive and ten negative residual clinical biospecimens collected by nasopharyngeal (n=7), oropharyngeal (n=1), or NP/OP combination (n=12) swabs into universal transport medium. Each sample was split into three separate 100 µL aliquots and processed independently through Qiagen (QIA), Macherey Nagel (MN), and bioMérieux easyMAG (EMAG) RNA extraction kits with 100 µL elution volumes, yielding a total of 60 extraction samples. All 30 positive and all 30 negative extraction samples were correctly assigned by the assay in comparison to orthogonal results from the outside laboratory (Table 6). Furthermore, the Ct values for each unique clinical sample were similar across the three different extraction methods assessed (Supplemental Table 1).

To calculate the confidence interval for the analytic accuracy of the assay, we counted every unique extraction of a clinical sample or of a synthetic positive extraction control once. We also counted each biologically unique nucleic acid source once. Based on this accounting, we had 47 true positive and 33 true negative results (Table 7), yielding 100% analytical sensitivity (95% CI: 0.9244-1) and 100% analytical specificity (95% CI: 0.8957-1).

**Table 7:** Analytical Accuracy

| **Sample Type** | **True Positive** | **True Negative** | **False Positive** | **False Negative** |
| --- | --- | --- | --- | --- |
| *Clinical Samples* | 30 | 30 | 0 | 0 |
| *Extraction Controls* | 13 | 0 | 0 | 0 |
| *Nucleic Acid* | 4 | 3 | 0 | 0 |
| *Total* | 47 | 33 | 0 | 0 |

### Limit of Detection

We utilized the synthetic positive extraction control and a synthetic negative extraction control for validation of the lower limit of detection for the assay. These extraction controls are delivered in a synthetic transport matrix including human gDNA. We diluted 200 viral cp/µL standard control into the negative extraction control matrix containing 75 human gDNA copies/µL to achieve final viral targets of 20, 10, and 5 copies per microliter. We performed 2 to 4 extractions of aliquots at each copy number target using a combination of Qiagen and Macherey Nagel extraction kits; 2 to 5 PCR replicates were performed for each unique extraction. At 5 viral copies/µL input to extraction, we successfully detected N1 and N2 in 20 of 20 replicates (Table 8), establishing a reliable limit of detection (LOD). The average N1 Ct at this LOD was 36.0 for the Qiagen extraction and 36.7 for the Macherey Nagel extraction; the average N2 Ct values were 35.8 and 37.2 respectively. Overall, the experiment had 97% accuracy at 1X to 2X of the limit of detection (5-10 cp/µL). One of ten PCR replicates at the 10 cp/µL input level returned an inconclusive result, detecting N1 but not N2.

**Table 8:**
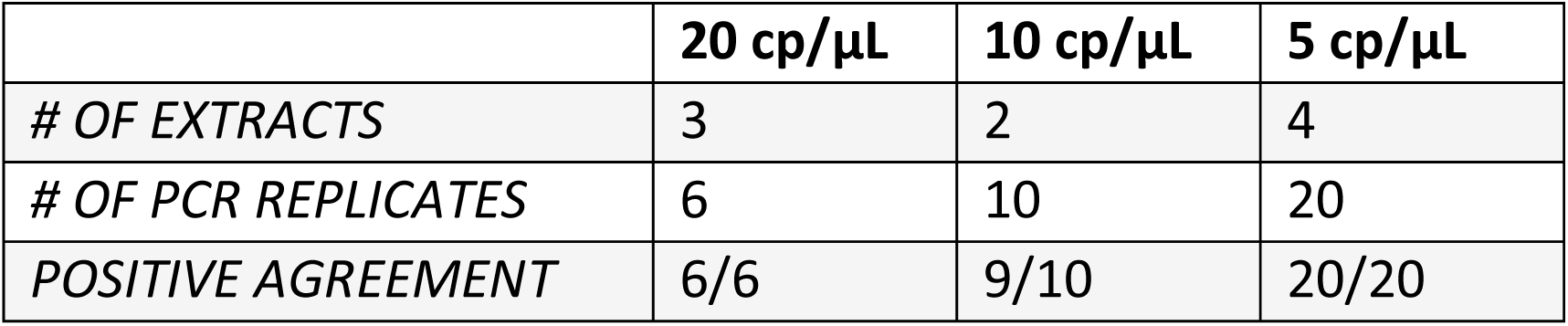
Limit of Detection

### Analytical Precision

Reproducibility of results between RNA replicates of both clinical samples and synthetic or genomic isolate controls was characterized across these validation experiments. We tested a total of 24 replicate sample pairs, not including the limit of detection replicates described above. Comparing across separate PCR plates (inter-run), we had complete concordance between 4 clinical sample pairs and 6 control sample pairs (Table 9). Within single PCR plates (intra-run), we showed 100% concordance between 6 clinical sample and 8 control sample pairs.

**Table 9:** Analytical Precision

|  | **Concordant Inter-run Comparisons** | **Concordant Intra-run Comparisons** |
| --- | --- | --- |
| *Positive Clinical Replicate Pairs* | 2/2 | 3/3 |
| *Negative Clinical Replicate Pairs* | 2/2 | 3/3 |
| *Positive Synthetic/Isolate Replicate Pairs* | 4/4 | 6/6 |
| *Negative Synthetic/Isolate Replicate Pairs* | 2/2 | 2/2 |

## Discussion

Using a combination of Emergency Use Authorized (EUA) and research use only (RUO) reagents, our multi-disciplinary clinical science team was able to validate analytic performance of this SARS-CoV2 detection assay in a very short time frame (5 days) and in the face of extraordinary barriers to the acquisition of EUA-cleared reagents. Our results demonstrate robust performance of a 10 µL volume, 384-well format qRT-PCR on the QuantStudio 5 instrument using a RUO Promega one-step mastermix. This format will support test throughput and increase the number of reactions possible in the current landscape of scarce reagents. Our results also demonstrate suitable performance of the Macherey-Nagel Nucelospin Virus extraction kit in comparison to two other EUA methods. We developed robust biosafety specimen handling procedures to ensure safety of all laboratory staff and a custom informatics system to promote scalability of sample accessioning, testing, and resulting.

Given the immense scale of the COVID-19 pandemic, it is predicted that reagent shortages for testing will be a continuous problem [6]. The ability of high-complexity, College of American Pathologist/CLIA88 compliant laboratories to rapidly and reliably adapt locally developed and validated testing procedures will play a key role in the United States’ ability to confront this clinical need. Other groups have recently provided important data on the relative performance of different primer/probe sets for SARS-CoV2 [4,7]. Intriguingly, Bruce et al. [8] provide data indicating that qRT-PCR detection of SARS-CoV2 could be successful on nasopharyngeal swab samples without any prior RNA extraction, at least as a screening mechanism. Innovative, rapid, and robust development and validation of new laboratory developed procedures will be required to help address the COVID-19 pandemic. In this setting, our work provides a blueprint for rapid characterization of new assay components and approaches as future supply chain shortages emerge.

## Acknowledgements

The authors wish to thank the research community of the University of Minnesota at-large who stepped-up in a time of both critical need and short supply to donate extraction materials, personal protective equipment, laboratory space, and encouragement to this this effort. We thank Drs. Jakub Tolar, Timothy Schacker, Leo Furcht, and Anthony Killeen for their support of this project and Drs. Ashley Haase, Stephen Rice, and Wade Bresnahan for donating their laboratory spaces to support this work. We thank Dr. Sara Vetter (Minnesota Department of Health) and Drs. Sophie Arbefeville and Patricia Ferrieri (University of Minnesota) for arranging access to critical clinical samples and discussion on study design. We thank Shea Anderson, Dinesha Walek, Karina Sartorio, and Cody Hoffman (University of Minnesota Genomics Center) and Shannon Gascoigne and Ann Fickle (M Health Fairview) for logistical and operational support. We thank Julie Jarvis and Aaron Beckman for their extraordinary effort to transport critical reagents from manufacturing facilities to our laboratory. ACN would like to thank Dr. Michelle Doyle for her critical feedback and support.

## TABLES AND FIGURES

**Supplemental Table 1:** Ct Values for Clinical Samples on Three Extraction Methods

| Sample | N1 | N2 | RP | Expected | Result |
| --- | --- | --- | --- | --- | --- |
| POS1_EMAG | 25.41 | 25.50 | 24.39 | + | Positive 2019-nCoV |
| POS1_MN | 25.70 | 26.19 | 28.83 | + | Positive 2019-nCoV |
| POS1_QIA | 25.04 | 25.98 | 28.50 | + | Positive 2019-nCoV |
| POS2_EMAG | 21.13 | 21.27 | 31.28 | + | Positive 2019-nCoV |
| POS2_MN | 21.56 | 21.99 | 32.10 | + | Positive 2019-nCoV |
| POS2_QIA | 20.75 | 21.69 | 31.80 | + | Positive 2019-nCoV |
| POS3_EMAG | 30.93 | 30.96 | 31.95 | + | Positive 2019-nCoV |
| POS3_MN | 32.06 | 33.20 | 33.12 | + | Positive 2019-nCoV |
| POS3_QIA | 30.06 | 31.97 | 31.23 | + | Positive 2019-nCoV |
| POS4_EMAG | 26.57 | 26.84 | 26.97 | + | Positive 2019-nCoV |
| POS4_MN | 26.15 | 26.94 | 26.26 | + | Positive 2019-nCoV |
| POS4_QIA | 26.35 | 27.01 | 27.80 | + | Positive 2019-nCoV |
| POS5_EMAG | 29.45 | 30.00 | 27.38 | + | Positive 2019-nCoV |
| POS5_MN | 30.67 | 31.79 | 29.13 | + | Positive 2019-nCoV |
| POS5_QIA | 28.00 | 28.61 | 26.94 | + | Positive 2019-nCoV |
| POS6_EMAG | 24.06 | 24.34 | 30.51 | + | Positive 2019-nCoV |
| POS6_MN | 25.40 | 26.00 | 33.81 | + | Positive 2019-nCoV |
| POS6_QIA | 29.17 | 29.95 | 35.29 | + | Positive 2019-nCoV |
| POS7_EMAG | 20.84 | 20.97 | 28.27 | + | Positive 2019-nCoV |
| POS7_MN | 21.47 | 22.13 | 29.90 | + | Positive 2019-nCoV |
| POS7_QIA | 19.82 | 21.29 | 27.97 | + | Positive 2019-nCoV |
| POS8_EMAG | 18.51 | 18.47 | 27.47 | + | Positive 2019-nCoV |
| POS8_MN | 18.78 | 19.08 | 27.74 | + | Positive 2019-nCoV |
| POS8_QIA | 17.58 | 18.78 | 26.93 | + | Positive 2019-nCoV |
| POS9_EMAG | 27.54 | 27.78 | 28.99 | + | Positive 2019-nCoV |
| POS9_MN | 28.20 | 28.97 | 30.18 | + | Positive 2019-nCoV |
| POS9_QIA | 26.54 | 27.68 | 28.80 | + | Positive 2019-nCoV |
| POS10_EMAG | 23.21 | 23.44 | 28.40 | + | Positive 2019-nCoV |
| POS10_MN | 23.59 | 24.53 | 30.72 | + | Positive 2019-nCoV |
| POS10_QIA | 23.32 | 24.32 | 29.76 | + | Positive 2019-nCoV |
| NEG1_EMAG |  |  | 28.71 | - | Not Detected |
| NEG1_MN |  |  | 29.79 | - | Not Detected |
| NEG1_QIA |  |  | 30.18 | - | Not Detected |
| NEG2_EMAG |  |  | 27.32 | - | Not Detected |
| NEG2_MN |  |  | 28.03 | - | Not Detected |
| NEG2_QIA |  |  | 27.44 | - | Not Detected |
| NEG3_EMAG |  |  | 28.38 | - | Not Detected |
| NEG3_MN |  |  | 27.60 | - | Not Detected |
| NEG3_QIA |  |  | 28.23 | - | Not Detected |
| NEG4_EMAG |  |  | 25.47 | - | Not Detected |
| NEG4_MN |  |  | 26.05 | - | Not Detected |
| NEG4_QIA |  |  | 25.14 | - | Not Detected |
| NEG5_EMAG |  |  | 28.40 | - | Not Detected |
| NEG5_MN |  |  | 29.90 | - | Not Detected |
| NEG5_QIA |  |  | 28.66 | - | Not Detected |
| NEG6_EMAG |  |  | 28.41 | - | Not Detected |
| NEG6_MN |  |  | 29.99 | - | Not Detected |
| NEG6_QIA |  |  | 28.31 | - | Not Detected |
| NEG7_EMAG |  |  | 27.52 | - | Not Detected |
| NEG7_MN |  |  | 29.62 | - | Not Detected |
| NEG7_QIA |  |  | 27.28 | - | Not Detected |
| NEG8_EMAG |  |  | 28.97 | - | Not Detected |
| NEG8_MN |  |  | 30.78 | - | Not Detected |
| NEG8_QIA |  |  | 28.57 | - | Not Detected |
| NEG9_EMAG |  |  | 30.76 | - | Not Detected |
| NEG9_MN |  |  | 32.55 | - | Not Detected |
| NEG9_QIA |  |  | 29.97 | - | Not Detected |
| NEG10_EMAG |  |  | 26.76 | - | Not Detected |
| NEG10_MN |  |  | 31.11 | - | Not Detected |
| NEG10_QIA |  |  | 27.98 | - | Not Detected |

EMAG: easyMAG; MN: Macherey Nagel; QIA: Qiagen

